# Accuracy and Precision of Social Relationship Indices

**DOI:** 10.1101/2021.04.25.441321

**Authors:** Alexander Mielke, Liran Samuni

## Abstract

Combining interaction rates of different social behaviours into social relationship indices to represent the structure of dyadic relationships on one underlying dimension is common practice in animal sociality studies. However, the properties of these relationship indices are not well explored – mainly because, for real-world social systems, the ‘true’ value of relationships is unobservable. Here, we use simulation studies to estimate the accuracy and precision of three relationship indices: the Dyadic Composite Sociality Index, the Composite Relationship Index, and the Dynamic Dyadic Sociality Index. We simulated one year of social interactions for multiple groups of 25 individuals and 4 interaction types with different properties, and tested the impact of different focal follow regimes, data densities and sampling conditions on the representation of social relationships. Accuracy and precision of social relationship indices were strongly driven by sample size, similar to simple interaction rates. Under the assumption that there was a clear, one-dimensional relationship underlying interactions, and that different interaction types constituting an index were highly correlated, indices indeed increased accuracy over single interaction rates for small sample sizes. Including uninformative constituting behaviours (i.e., those not highly correlated with the underlying relationship dimension) reduced the accuracy of all indices. The precision of each index (i.e., whether multiple simulated focal follow regimes achieve the same dyadic values for the same data) was generally poor and was driven by the precision of the least precise constituting behaviour, making them less precise than some single interaction rates. Our results showed that social relationship indices do not remove the need to have sufficient data for each individual constituting interaction type. Index quality was defined by the least accurate and precise constituting interaction type. Indices might only be useful if all constituting interaction rates are highly correlated and if there are clear indications that one dimension is sufficient to represent social relationships in a group.

## Introduction

Animals in stable social groups maintain a variety of differentiated social relationships with group members, from kin relations and mating partners, to cooperative social ties between non-kin (Cheney, 2011; Gero et al., 2009; Kern & Radford, 2016; Mitani, 2009; Riehl & Strong, 2018; Samuni et al., 2021; Schülke et al., 2010; Silk et al., 2003; Wilkinson et al., 2016). Individuals’ social environment appears crucial for improved development, social status, health, stress management, reproductive success, and survival (Cameron et al., 2009; Riehl & Strong, 2018; Silk et al., 2010b; Wittig et al., 2016). As such, a large body of research is dedicated to studying the contribution of differentiated social relationships to fitness consequences (Snyder-Mackler et al., 2020). Equally, much research has been devoted to the cognitive challenges arising from differentiated and changing social relationships and their role in competition for resources, mates, and cooperation partners (Seyfarth & Cheney, 2015). Relying on the characterisation of ‘social relationships’ to ask questions about social complexity, cooperation, and fitness makes it vital to quantify the underlying distribution and strength of dyadic relationships accurately. However, data collection in the behavioural sciences is messy and patchy as only some animals can be followed every day. We often do not know what an ‘accurate’ description of relationships would look like, as we do not know how individuals represent their connections with others. It is therefore important to explore the assumptions and properties of our social relationship indices, to enable researcher to make the best decisions about how to describe relationships within their study group.

To calculate social relationship indices, researchers often use behavioural observations of species-typical dyadic interactions (e.g., grooming, coalition formation, spatial proximity) aggregated over time, and standardised for observation effort and/or individual gregariousness and/or group-typical interaction rates (Silk et al., 2013). These (standardised) interaction rates are combined to create an index that represents the quality of the underlying relationship of each dyad on a single dimension. This seemingly straightforward application makes indices popular. However, if not applied correctly, sociality indices affect the validity and robustness of results and limit replicability (Whitehead, 2008). Attempts to validate the reliability and robustness of interaction-based indices are largely absent – more detailed attempts exist for association indices (Davis et al., 2018; Whitehead, 2008). Importantly, researchers face a number of choices when creating their study-specific relationship index, posing problems for the replicability of researcher degrees of freedom across studies (Wicherts et al., 2016). Researchers have to identify interaction types to include in their index, how to standardise them, how to weight them, and how to combine them (Silk et al., 2013). Another difficulty is sample size: fewer data points come with increased measurement error and decreased power and precision when making dyadic indices (Whitehead, 2008). Many interaction types, such as food sharing or coalitionary support, are rare in wild animals given the number of individuals in a group, making the calculation of interaction rates unreliable (Mielke et al., 2021), but it is currently unknown whether combining several interaction types with independent error rates increases or decreases overall error.

A variety of social relationship indices is currently used across species and studies. Here, we focus on three indices: the Dyadic Composite Sociality Index (CSI or sometimes DSI; Sapolsky et al., 1997; Silk et al., 2003), the Composite Relationship Index (CRI; Crockford et al., 2012), and the Dynamic Dyadic Sociality Index (DDSI, Kulik, 2015). Each of these indices makes different assumptions (please see SI for detailed descriptions of index formulas and underlying assumptions), but their main goal is to describe social relationships on a single dimension. The CSI and CRI aggregate multiple interaction rates and quantify how dyads compare with average rates within their group. The CSI only includes socio-positive interaction, while the CRI also incorporates socio-negative interactions and can weigh rare interactions more strongly. The DDSI can include socio-positive and socio-negative interaction types and has dynamic features resembling the Elo rank index (Albers & De Vries, 2001): positive interactions increase dyadic values from that day onwards, while negative interactions decrease values (De Moor et al., 2020; Mielke et al., 2017; Samuni et al., 2018). The impact of each interaction type is frequency-dependent, with rare interaction types weighing more strongly; however, weights can be set manually.

Relationship indices make assumptions that are rarely met or validated, some of which have the potential to increase inter-individual measurement error and result in outcome uncertainty. On the most basic level, we assume that social relationships can be presented on a single dimensions without losing important information – this is by no means a given and not inherent in the original understanding of ‘social relationships’ (Hinde, 1976). We assume that the index we choose is accurate (approaches the ‘real’ relationship value) and precise (multiple applications of different data for the same timeframe should converge on a similar solution). The approaches should be reproducible – different research groups with the same dataset should independently converge on the same results (LeBel et al., 2018). We also make numerous assumptions regarding the data used to compile the indices. We assume that our sample size is *sufficient* to construct accurate social tendencies, that the types of social interactions included are a *meaningful* representation of differentiated social relationships, or that our dataset represents balanced sampling effort of individuals and interaction types (or instead that the indices used are robust against unbalanced sampling efforts). All those assumptions are not method-specific and should be considered independently of the choice of method.

Method-specific assumptions likely also increase measurement error and uncertainty. Being composite measures, relationship indices incorporate assumptions regarding the relative contribution of various interaction types to the overall measure. For example, the CSI assigns the same importance to rare and common affiliative interaction types (e.g., body contact and food sharing), even though these might reflect relationships differently and might have different measurement error. The CSI and CRI are variance-dependent, so dyadic values can be high either because the dyad interacts a lot, because other dyads interact little, or because too few data are available – the interaction type with the largest number of zeros will potentially have an outsized influence on the final index. When rates are standardised against the group mean, individual differences in interaction rates are ignored – ‘friendship’ might manifest itself very differently across individuals, reducing the ability of an index to predict observed behaviour. Further, social indices treat ‘social relationships’ as either fixed averaged parameters (CSI and CRI) or dynamic sequential processes (DDSI), but we have not yet validated which approach is more biologically meaningful, or how varying sampling effort and timing in combination within each approach may affect indices values. All indices press the assumed underlying relationships into a specific distribution (right-skewed with a mean of 1 for the CSI, normally distributed with a mean of 0 for the CRI, normally distributed with a mean of 0.5 for the DDSI) – independent of the actual distribution of interactions in the social group.

Despite the potential for spurious inferences if assumptions are violated, attempts to validate existing indices are largely lacking. Consequently, evaluating how the different indices perform when assumptions are not met, or assessing which index provides a more meaningful representation of ‘true’ occurrence, is a crucial next step in advancing the reliability and replicability of social relationship studies. As a first step to increase robustness of social relationship assessments, we must identify the set of tools that allows us to minimize uncertainty and measurement error in existing methods. Although we operate under the premise that an underlying distribution of social relationships exists within our data, we do not know what the ‘real’ underlying distribution is, making validation attempts challenging based on real-world data. To overcome this challenge, in this study, we simulated datasets of social groups of animals, with each group member ‘followed’ daily, thus representing an optimal social interaction dataset – both the underlying relationship dimension and the ‘full’ dataset are known. We conducted simulations according to a defined set of rules regulating the underlying distribution of different types of interaction. Each dyad was assigned a specific ‘expected probability’ of affiliative interaction (see Supplementary for description of simulation process), while different ‘interaction types’ skew the distribution to be more or less selective. In more selective interaction types, only dyads with high probability of affiliation would interact, while in less selective interaction types, dyads with low probabilities also have a chance to interact (Whitehead, 2008). Interaction probabilities of socio-negative interaction types followed adapted rules that meant they were negatively correlated with the socio-positive interaction types, as typically so in real-world data. The underlying ‘expected probabilities’ serve as the ‘true’ basis of the social environment in the group, upon which different applications of social relationship indices can be compared and validated.

Using the simulated dataset, we focused our questions on both intra- and inter-index validation steps. We assessed the quality of each index by varying the a) overall sampling effort, b) balance of sampling effort across subjects, and c) measurement error, by incorporating interaction types that diverge from the ‘true’ relationship and thus create random variation. We then compared how the different indices performed under these scenarios. We compared the performance of indices 1) against the ‘true’ interaction probabilities and 2) against the indices calculated based on the full datasets, which incorporates all interactions for all individuals of the group. These two measures represent the accuracy of the index. We also 3) determine the impact of sampling variation – if taking data from the same overall datasets but with different focal individuals chosen on different days, how similar are the resulting indices?

## Methods

### Generating Data

To assess the robustness of the different social relationship indices, we simulated social interactions between individuals within ten different social groups of 25 individuals (equals 300 dyads) over a one-year period. For each individual, we generated a daily dataset of social interactions during 12 observation hours. Each dyad was randomly assigned an interaction probability, and probabilities were subsequently standardised within individuals to sum to 1 to create the expected interaction probabilities from one partner to the other – see Supplementary for details and GitHub for scripts.

The expected interaction probabilities of individuals with specific partners represent their ‘social relationship’ with this individual: dyads with high expected probabilities interact positively a lot, while those with low expected probabilities would hardly interact positively with each other but show a higher level of aggression. Socio-positive interaction probabilities are roughly symmetrical within dyads but right-skewed overall, so few dyads have a high probability to interact a lot (see Fig. 1). Socio-negative interaction probabilities were created by repeatedly squaring the expected interaction probability and subtracting it from 1, and then standardising it to sum to 1 within individuals. This created a right-skewed distribution that was negatively correlated with the expected interaction probabilities and the socio-positive interaction distributions. Random error is added by making each interaction choice based on a random sample of all group members – simulating flexible spatial patterns in social animals. Therefore, both the expected probabilities and partner availability defined who would interact in case interactions occur.

**Figure 1:**
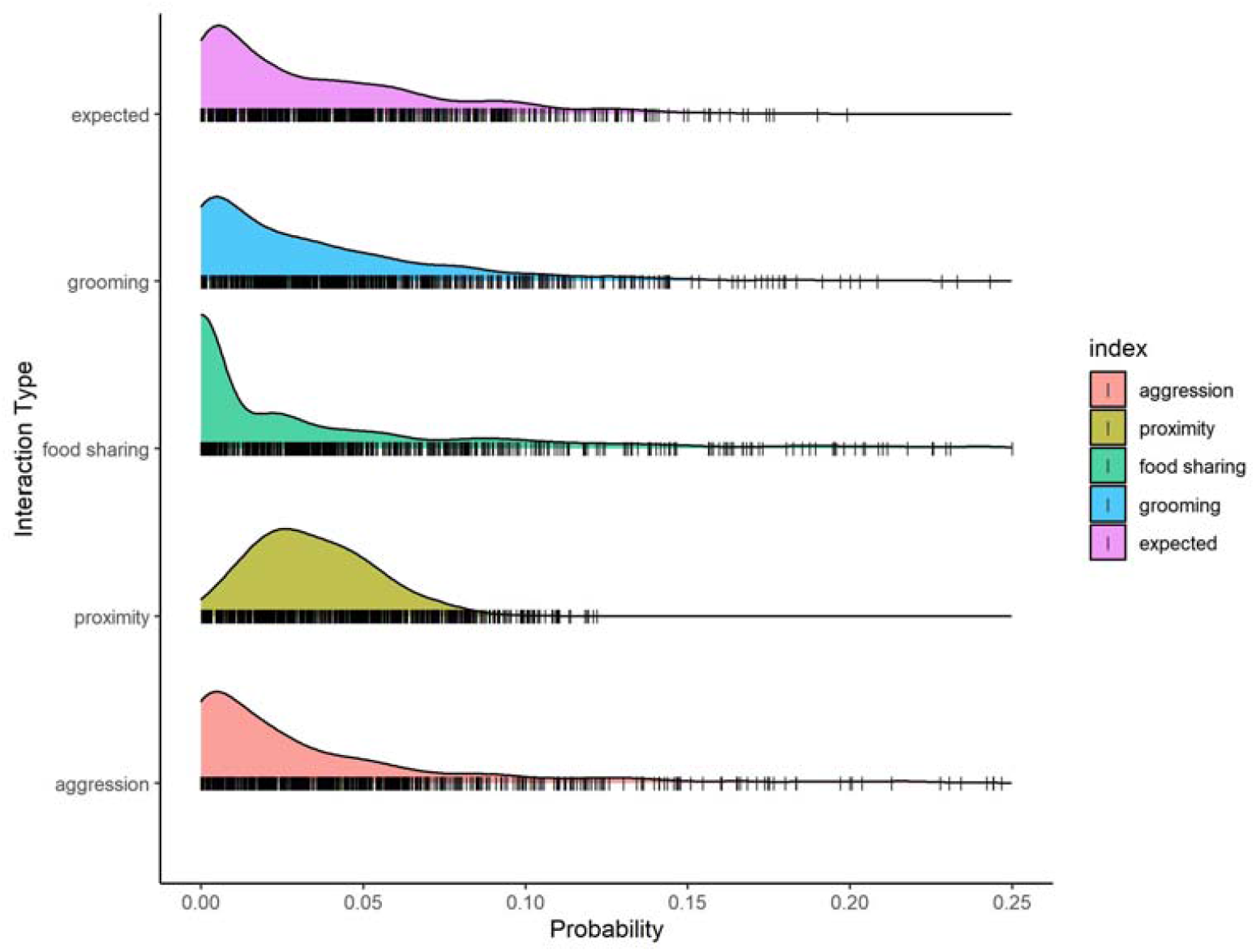
Density of interaction distributions for each of the four simulated interaction types and the expected probability. High selectivity (e.g., for food sharing) creates stronger right skew, while less selectivity (e.g., proximity) leads to more even distributions

We aimed to simulate interactions types that are typically used when constructing social relationship indices. For each individual and day, a fixed number of interactions of each type was set. As in real social systems, the frequency and selectivity of occurrence of interaction types varied. The frequency value of the behaviour represents the average number of interactions of that type exhibited per day. The selectivity of the behaviour, reflected in higher or lower coefficient of variance for different interaction types, signifies partner choice fidelity for the respective interaction type. If selectivity is high, only a small number of partners are ever chosen, and the interaction distribution is highly right-skewed; if selectivity is low, the likelihood of choosing each partner is roughly even. A behaviour of low selectivity has low explanatory power of the relationship value and vice versa. Overall, we simulated three socio-positive and one socio-negative interaction types, with varying frequencies and certainties – their distributions can be seen in Figure 1:

1. Behaviour 1 (“grooming”) – socio-positive behaviour of medium frequency and selectiveness. We simulated a frequency of 3 interactions per day with medium selectivity (coefficient of variance ∼ 1.2).
2. Behaviour 2 (“food sharing”) - socio-positive behaviour of low frequency and high selectiveness. We simulated a frequency of 1 interaction per day with high selectivity (coefficient of variance ∼ 1.8).
3. Behaviour 3 (“spatial proximity”) – a common socio-positive behaviour of low selectiveness. We simulated a frequency of 8 interactions per day with low selectivity (coefficient of variance ∼ 0.8).
4. Behaviour 4 (“aggression”) - socio-negative behaviour of medium frequency and selectiveness. We simulated a frequency of 3 interactions per day with medium selectivity (coefficient of variance ∼ 1.2).

Unless otherwise indicated, ‘CSI’ refers to the Dyadic Composite Sociality Index using behaviours 1-3, ‘CRI’ refers to the Composite Relationship Index using all four behaviours (with Behaviour 2 weighted stronger and Behaviour 4 having a negative impact), and the DDSI refers to the Dynamic Dyadic Sociality Index on the last day of the observation period, using all four behaviours. For all indices, we test the impact of ‘bad’ or uninformative behaviours by exchanging Behaviour 1 (medium frequency and selectivity) with an unrelated behaviour.

### Simulations

We conducted a series of simulations to investigate the measurement accuracy of the three indices (i.e., CSI, CRI and DDSI) and rates of single interaction types. We generated the complete datasets, with every interaction for every individual known for the entire period. Each individual had 3 grooming bouts, 1 food sharing event, 8 proximity events, and 3 aggression events per day – the total number of interactions for the whole group therefore amounted to about 27,000 grooming and aggression events, 9,000 food sharing events, and 72,000 proximity events. These constitute our ‘optimal’ value and serve as a test for the accuracy and precision of the simulations. One focal individual was randomly chosen per day, and the number of observation days was varied randomly. After, we included a number of conditions (differing observation effort, sampling bias, inclusion/exclusion of specific interaction types; see below) to test the accuracy and precision of the indices in different circumstances. It is worth noting that our datasets contain less noise than real-world data and are therefore expected to show clearer patterns with less data available. For example, in most primate species, grooming is not only reserved for relationship maintenance but also traded for infant access or use for reconciliation (Carne et al., 2011) or varies according to changes in the ecological or social environment (Brent et al., 2013), blurring the picture.

#### How are indices affected by the observation effort?

For each of the 10 generated datasets, we conducted 100 random selections of observation days, with one focal observed on each day (Mielke et al., 2021). We randomly selected between 30 and 360 observation days (360h – 4320h of observation, which is within the usual range for field studies), resulting in 10 x 100 datasets. We used the simulated datasets to evaluate the performance of the different indices for varying sampling efforts and under different conditions (i.e., sampling bias, inclusion/exclusion of specific interaction types; see below) to test the accuracy and precision of the indices.

For each condition, we report two main measures of index quality: *accuracy* and *precision*. We measured the *accuracy* of the index as the correlation between the observed index and a) the expected interaction probability, and b) the index based on the full dataset. We measured the precision as the variability of index values that dyads showed across simulations (Whitehead, 2008). We quantified the variability by simulating 100 random focal selections for the same number of observation days and calculating the 80%-interquantile ranges of the resulting indices –obtaining a range of values into which most of the dyadic index values fell. If this range is small, then sampling has little impact on dyadic values; if the range is large, following specific focal individuals on certain days skews the results, even for this artificially clean data set. For precision, we focus on two sample sizes as examples for high or low data density: 60 observation days (or 1800 interactions an individual is on average exposed to as sender or receiver) and 360 observation days (or 10,800 interactions an individual is on average exposed to). For all questions, we present graphs to report the results of the simulations: for the accuracy, we present graphs showing the observation effort on the x-axis and the correlation coefficient with the ‘true’ values on the y-axis. For the precision, we plot the ‘true’ relationship value (based on all interactions) of a dyad on the x-axis, and the 80%-interquantile range of dyadic values across 100 randomised focal follow regimes on the y-axis. Thus, the x-axis represents dyadic relationship strength, while the y-axis represents precision.

#### Do indices outperform single interaction rates?

One of the main aims of social relationship indices is to improve representation in case few data points are available – the assumption being that pooling several interaction rates creates more robust measures than a single interaction rate. Comparing different interaction types and indices directly, we test whether indices are more accurate and precise than the single interaction types constituting them. We also test whether social indices have a better explanatory power of the expected interaction probabilities over the separate interaction types, by fitting a series of Generalized Linear Models (GLM) with either the indices or the single interaction types as the predictors and the expected probabilities as response. Due to the proportional nature of the expected probabilities and their range between 0 and 1, we fitted the models with beta error distribution and logit link function using the *betareg* function of the identically named package (Zeileis et al., 2016). To quantify the relative performance of each model, we used a measure of model fit, the Akaike’s Information Criterion (AIC; Burnham et al., 2011). Lower AIC values indicate improved model fit. We then calculated the ΔAIC of each model as the difference from the lowest AIC value, which has a ΔAIC of zero. We centered our inference on ΔAIC (Burnham et al., 2011).

#### Do non-indicative interaction types bias indices?

How robust are indices if inappropriate behaviours are included in the calculation? For example, spatial proximity is a poor indicator of sooty mangabey social relationships (Mielke et al., 2020), but was previously included in social relationship indices for the species (Mielke et al., 2017) – would the index still be robust? We evaluated the relative robustness of index output by swapping some interaction types with others of identical frequencies and selectivity that are otherwise poor indicators of the interaction probabilities (i.e., ‘uninformative’ behaviours). We achieved this by borrowing interaction data of the same behaviour types across datasets (e.g., ‘positive medium behaviour’ from one dataset replaced ‘positive medium behaviour’ in another and was thus non-indicative of the expected interactions in that dataset, while otherwise showing the same properties). The correlation between the different behaviour types and the expected values are depicted in Figure S2.

#### Does unbalanced collection effort bias indices?

In addition to being incomplete, behavioural data collection is prone to collection effort biases between individuals, as not all individuals are sampled at the same frequencies. Therefore, to address whether uneven sampling densities between individuals may impact measurement accuracy of the three indices, we conducted our simulations while varying the sampling probabilities of individuals, using 0.2, 0.5 or 0.8 as the probability for an individual to be sampled. Thus, some individuals are on average 4 times as likely to be sampled as others, for whom most individual will be derived when they are non-focal interaction partners. We then investigated the influence of varying sampling efforts across individuals and dyads on index values.

## Results

### How are indices affected by observation effort?

One central point of this study is that, even in these highly predictable and stable data, low sampling effort leads to high uncertainty in all three relationships indices – negatively affecting their accuracy and precision. Figure 2 shows the correlation between each of the three indices – CSI, CRI, and DDSI – with the expected interaction probability and with the same index calculate over the full dataset. Indices based on smaller sample sizes are less accurate, regardless of the method. In our simulated datasets the ‘optimum’ is roughly achieved from ∼15 interactions per dyad, meaning that 4,500 interactions (15 interactions for 300 dyads in a group of 25 individuals) are needed to achieve adequate accuracy – a large number for any observational study. The indices do not differ in their trajectories, but the CRI correlated less highly with the underlying probability distribution than the other two indices.

**Figure 2:**
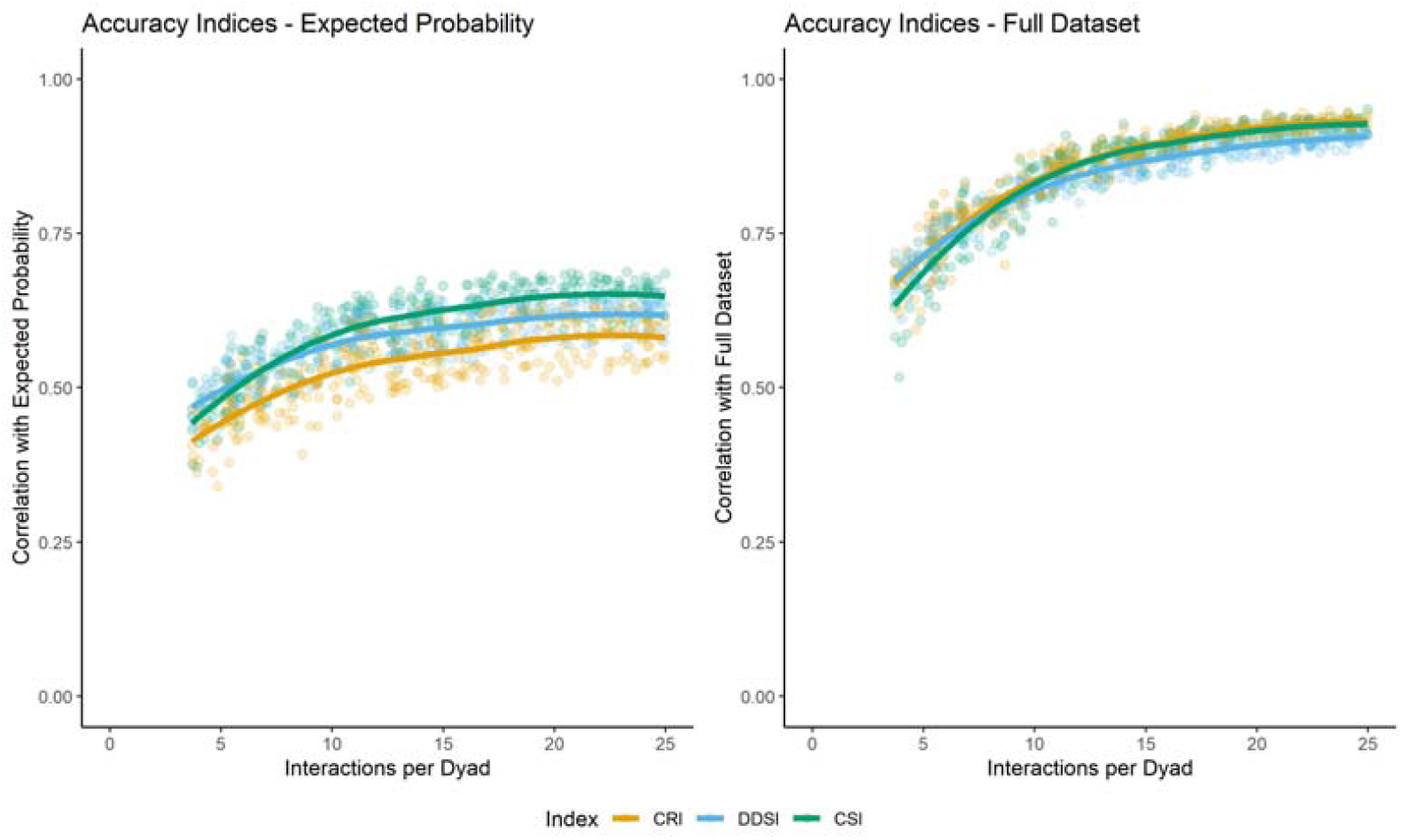
Pearson correlation of the CRI, DDSI, and CSI with the expected probability distribution (left) and the same index based on the full dataset (right). Each point marks one random selection of focal days.

For the precision of the indices, Figure 3 shows the dyadic value for the full dataset on the x-axis (standardised between 0 and 1 for interpretability), and on the y-axis it shows the 80%-interquantile range of values that a dyad was assigned with over the 100 randomized iterations of focal selections. A dyad with a value of 0.5 in the x-axis falls halfway between the maximum and minimum value of the index distribution, but if it has an interquantile range of 0.6 then index values ranging between 0.2 and 0.8 are likely – making dyadic values volatile and hard to interpret. Large interquantile ranges indicate high levels of imprecision. Our main assumption of focal follows is that the dyadic relationship values we find in our data are independent of the days on which individuals were followed. If a dyad sometimes falls into the top 20% of all dyads, and sometimes into the bottom 20%, that means that any result will be conditional on sampling biases. As Figure 3 shows, this is the case for all three indices, especially when data are sparse (i.e., 60 observation days or 1800 interactions). Note that for both the CRI and DDSI, dyads who never interact would have values around the centre of the distribution, leading to the observed shapes.

**Figure 3:**
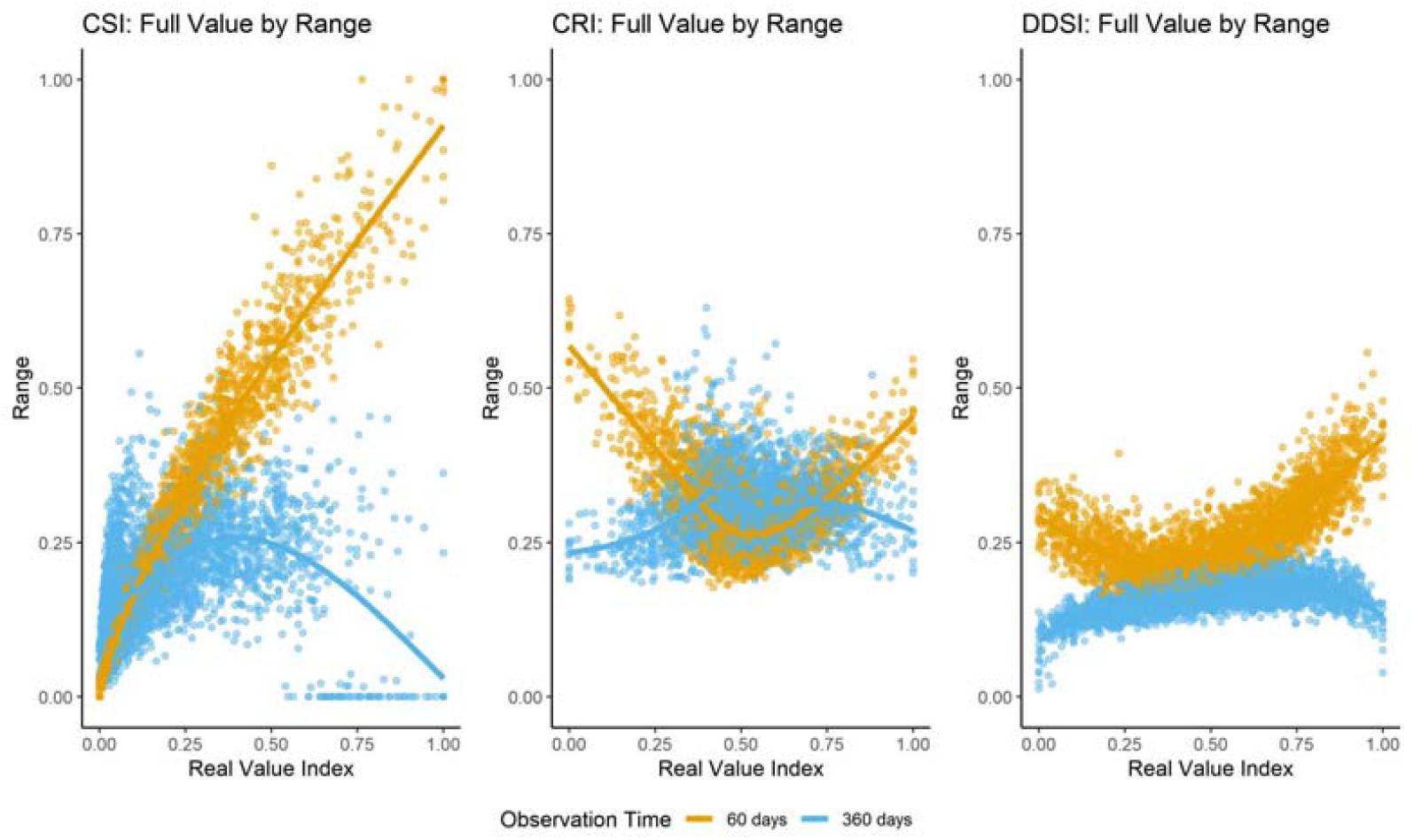
Precision for the three indices. The x-axis depicts dyadic values, standardised between 0 and 1 (i.e., 1 is the dyad with the highest value in the group). The y-axis depicts the 80%-interquantile range for 100 randomized sampling regimes. Colors depict different observation efforts. Higher range indicates lower precision. The differences in distribution shapes between indices are due to the fact that the CSI only includes socio-positive interaction types, so low values indicate no interactions (highly stable), while dyads without interactions are at the centre of the CRI distribution and have variable positions in the DDSI.

Smaller sample sizes (i.e., 60 observation days, around 1800 interactions in total) lead to more imprecision, especially for extreme values. For example, the dyad with the highest CSI value for the full dataset (1 on the x-axis) was regularly assigned the lowest and highest CSI values depending on the focal follow sampling scheme, leading to an interquantile range of 1. For the larger sample size (360 days), dyads with very high CSI values show higher consistency. For the CRI and DDSI, positive and negative extreme values exist (here standardized between 0 and 1), because dyads can exhibit more aggression than affiliation (Figure 3). For all three indices, with increasing data density, the uncertainty shifts from the extreme values to those in the centre of the distribution: while the indices correctly identify those dyads with extremely high and low values, the ones in between are poorly classified. For example, even with 360 observation days (around 10,800 interactions in total), a dyad with a ‘true’ standardized CSI value of 0.5 could be classified as 0.3 or as 0.7 – depending on which individual was sampled on which day. The assumptions of the indices lead to difference in the expression of precision, as dyads who never interact do not occupy the same space in the distribution for every sampling scheme. For the CSI, dyads who never interact always get a value of 0, so their assignment is precise – on the other hand, estimating the actual distribution of those dyads that do interact is highly dependent on the sampling. The CRI, on the other hand, and to a lesser degree the DDSI, suffer in precision across the distribution because negative interactions are included and shift the minimum value of the distribution.

### Do indices outperform single interaction rates?

One of the main reasons to apply a relationship index is to circumvent small sample sizes: if insufficient data are available for each individual interaction type, maybe combining them can reduce error. Figure 4 shows that this is indeed the case for small samples: as long as all included interaction types are highly correlated and measurement errors are uncorrelated (e.g., no seasonal effects exist), all relationship indices are able to improve accuracy of predictions of the underlying probability distribution compared to the individual interaction rates. The CRI does only do so for low sample sizes, and with increasing sample size, the advantage of the CRI index decreases – thus, once sufficient data is available for a single interaction type, that interaction type would, in theory, be a more accurate predictor than the CRI.

**Figure 4:**
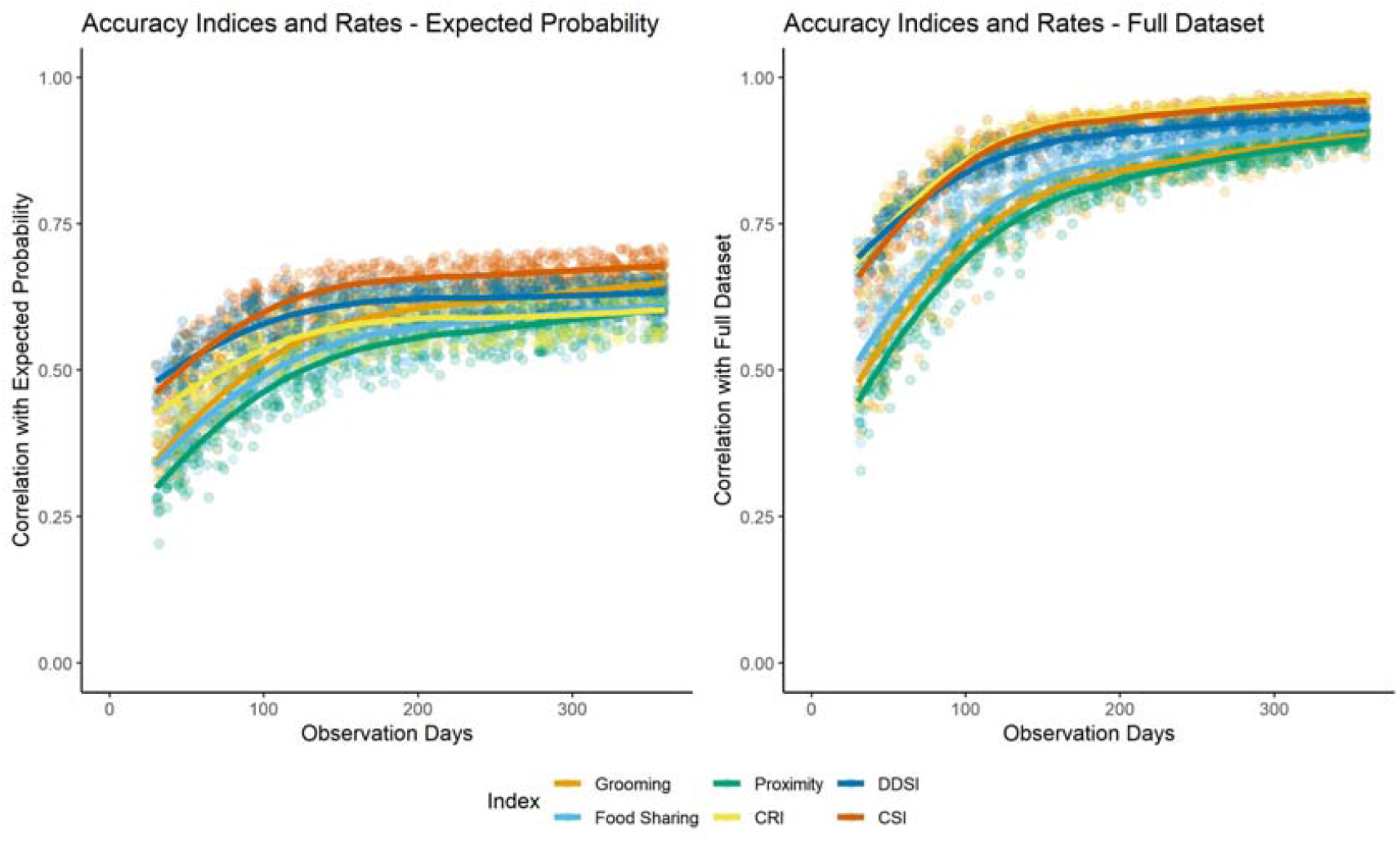
Pearson correlation between indices/rates and the expected probability and index/rate based on the full dataset. Comparison of accuracy for the three socio-positive interaction types and the three indices. With increasing sample size, the advantage of the indices is reduced.

In Figure 5, we observe the precision of the CSI compared to each of the socio-positive interaction types. Due to the different shapes of the DDSI and CRI distributions, which include negative interaction types, we omit them here, but results are the same. The CSI does not appear more precise than the individual interaction rates. It is also visible that the sample size (i.e., interaction frequency), which differs between the interaction types, impacts their precision, more so than the selectiveness of partner choice: ‘food sharing’, which had the highest selectivity but lowest frequency, is less precise than ‘proximity’ with the lowest selectivity but highest frequency. The precision of the CSI is most similar to the ‘food sharing’ rate, indicating that this relationship index is more strongly influenced by the most imprecise component.

**Figure 5:**
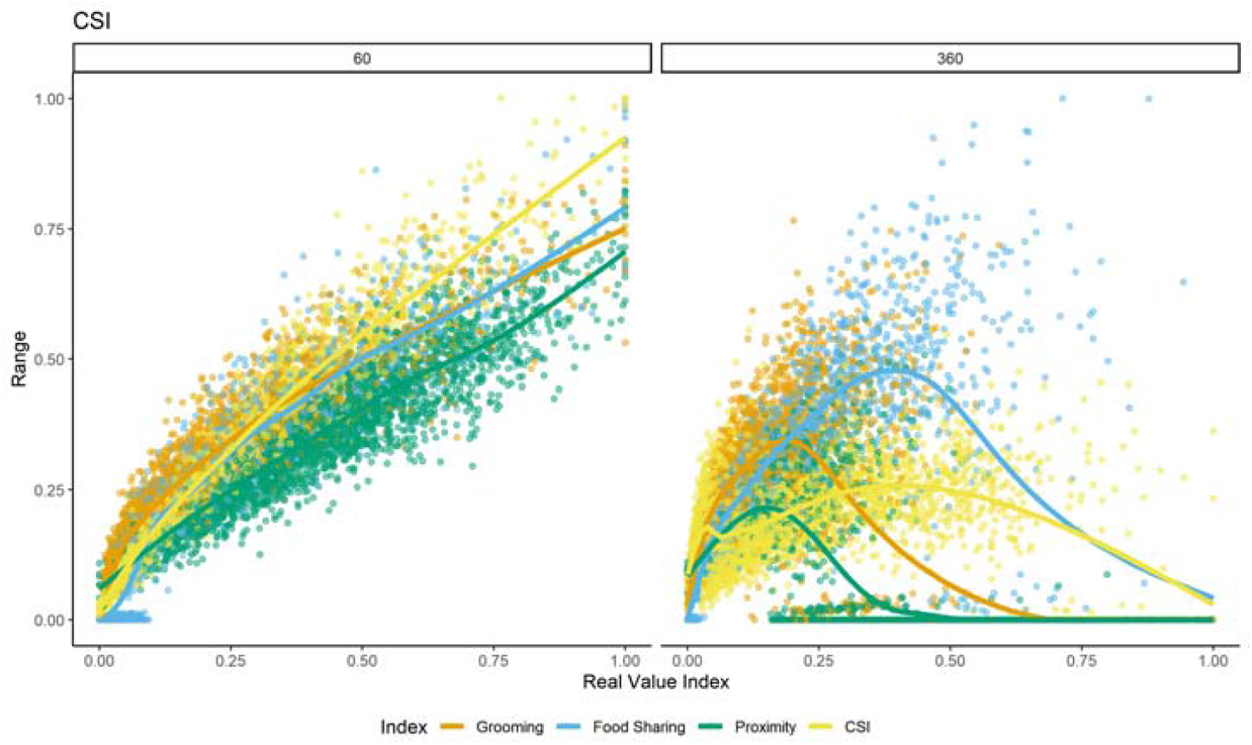
Precision for the CSI and three interaction rates. The x-axis depicts dyadic values, standardised between 0 and 1 (i.e., 1 is the dyad with the highest value in the group). The y-axis depicts the 80%-interquantile range for 100 randomized sampling regimes. On the left are randomized samples containing 60 observation days (around 1300 interactions), on the right 360 observation days (around 10,800 interactions). Higher range indicates lower precision.

As we can see in Figure 6, a CSI made without ‘food sharing’ (the least precise context) is more precise than one with all three socio-positive behaviours, while removing ‘proximity’ (the most precise context) reduces precision further as ‘food sharing’ has a stronger impact. Thus, index precision seems driven by the least precise constituting interaction type – which will usually be the rarest interaction type.

**Figure 6:**
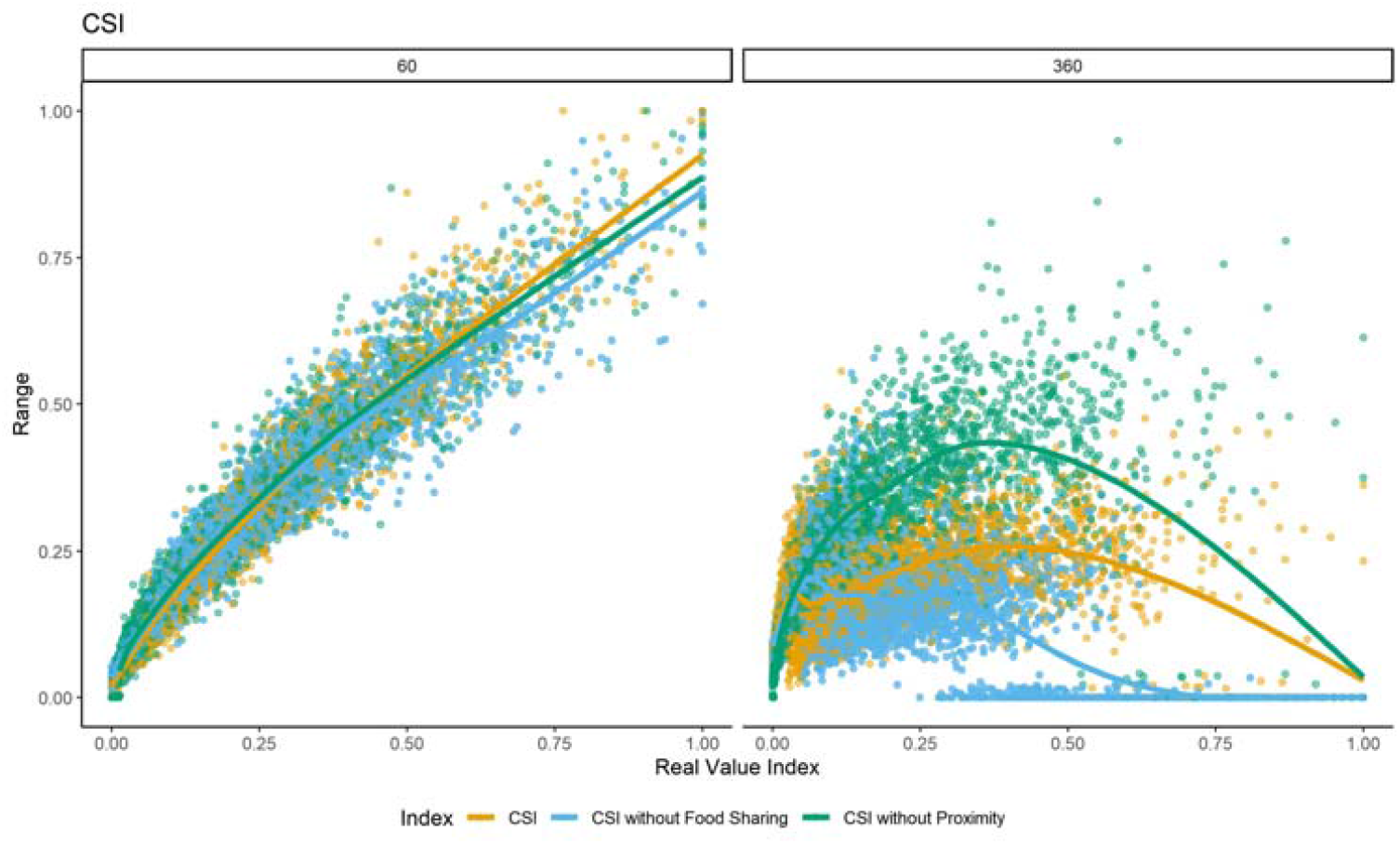
Precision for the CSI with and without the most precise (‘proximity’) and least precise (‘food sharing’) interaction type. The x-axis depicts dyadic values, standardised between 0 and 1 (i.e., 1 is the dyad with the highest value in the group). The y-axis depicts the 80%-interquantile range for 100 randomized sampling regimes. On the left are randomized samples containing 60 observation days (around 1300 interactions), on the right 360 observation days (around 10,800 interactions). Higher range indicates lower precision.

To test which of the social relationship indices or the constituting interaction rates best explain the expected interaction probabilities, we conducted beta regressions with the expected interaction probabilities as outcomes and calculated the ΔAIC values. Figure 7 shows the ΔAIC values obtained from four models containing either one of the three indices or all four interaction types as separate predictors. Including the interaction types as separate predictors in the same model captures a larger share of the variation in the expected probability distribution – the ΔAIC for the combination of independent rates is the best solution for most of the iterations, independent of sample size. Between the relationship indices, the DDSI, followed by the CRI, best explained the variation in the expected probability, with the CSI performing poorly in capturing the expected probability variation.

**Figure 7:**
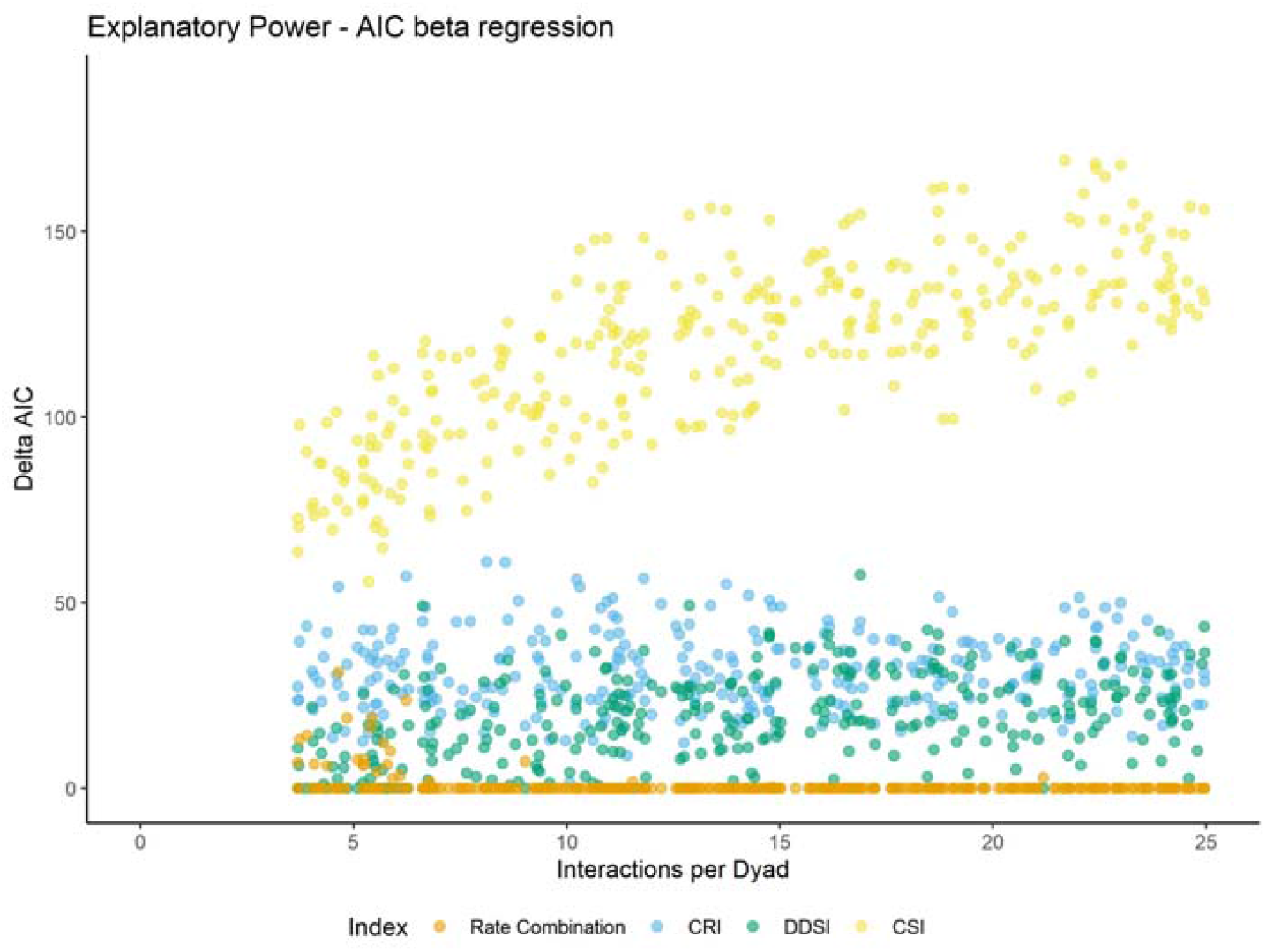
Delta AIC for the beta regression models with the expected probability as outcome and each index or the combination of interaction rates as predictor variables. Smaller values indicate more accurate models, with the combination of interaction rates as best model in most cases having a value of 0 throughout.

### Do uninformative interaction types bias indices?

Often researchers must decide whether to include interaction types into their relationship indices despite some uncertainty as to how well these reflect the underlying social relationship. However, we do not yet know whether or not this is a valid concern. Do we lose precision and accuracy by including an interaction type that does not represent the social relationship (i.e., uninformative) into the index? Figure 8 shows that all three indices are highly susceptive to uninformative (or ‘bad’) interaction types: in all indices, accuracy drops below the level of a single, informative, interaction type (i.e., an interaction correlated with the expected interaction probability). The effect is larger for the CSI, because it includes fewer components than the other two indices, each having a greater impact on its distribution. Figure 9 shows the difference for the performance of the CSI build with three informative behaviours, two informative and one uninformative behaviour, and one informative and one uninformative behaviour. As we can see, the fewer informative behaviours are involved, the stronger the negative impact of the uninformative behaviour is.

**Figure 8:**
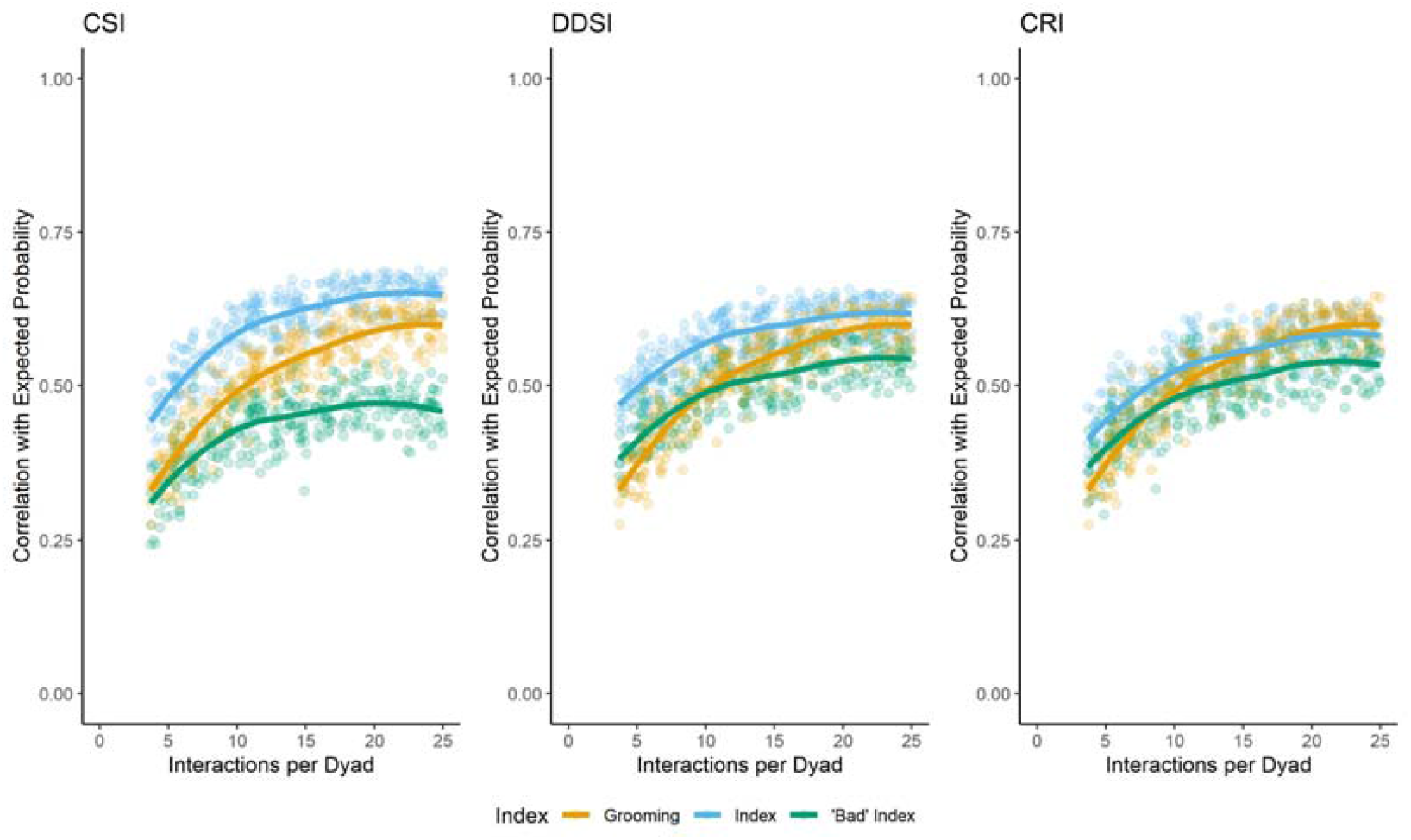
Pearson correlation between indices/rates and the expected probability. Comparison of accuracy for the three indices with and without an uninformative or ‘bad’ component. Interaction rate of Behaviour 1/’Grooming’ shown for comparison. Adding uninformative behaviours reduces accuracy dramatically.

**Figure 9:**
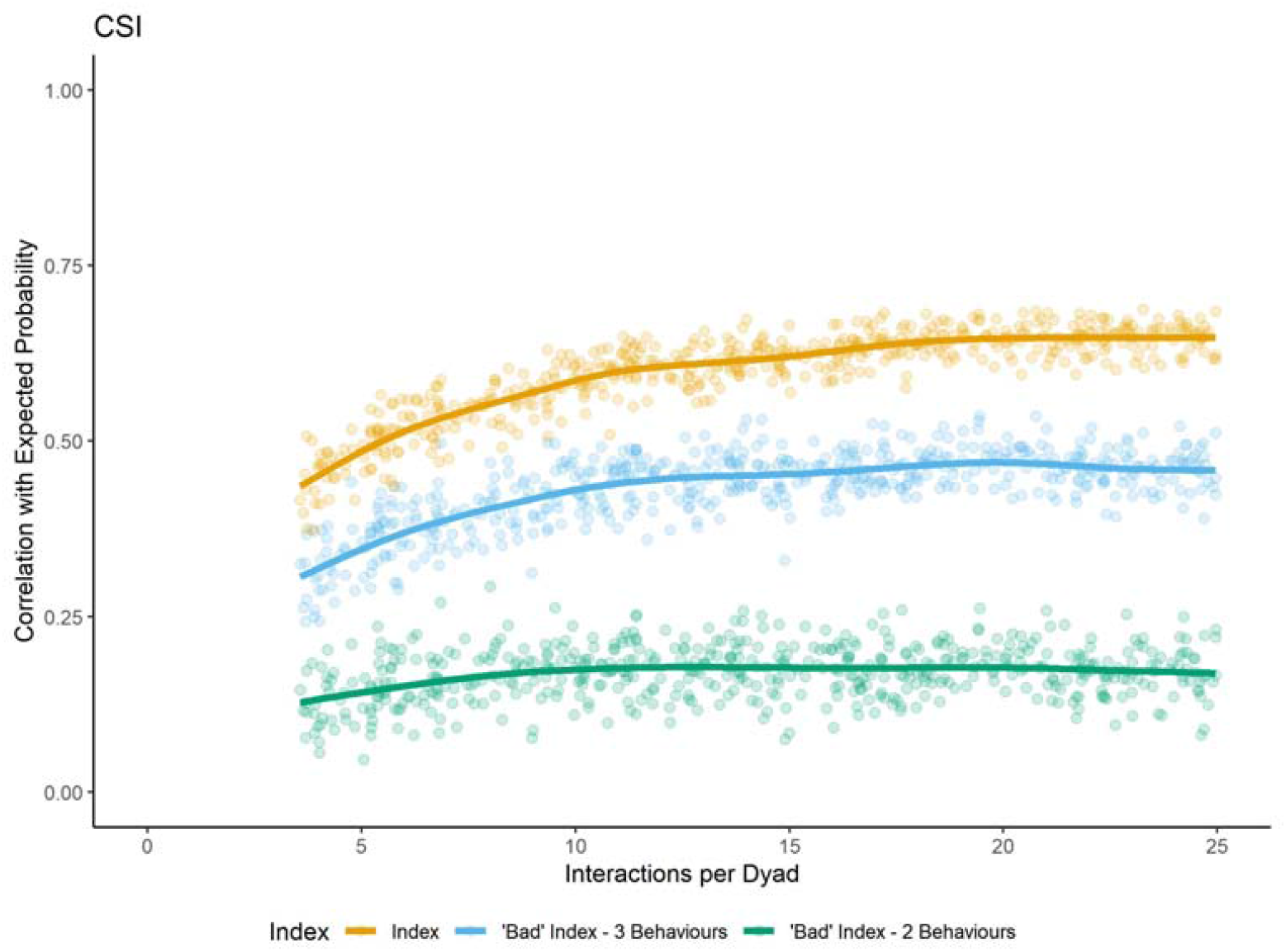
Pearson correlation between indices/rates and the expected probability. Comparison of accuracy for the CSI based on three informative, two informative and one uninformative (‘bad index’ – 3 behaviours), and one informative and one uninformative behaviours (‘bad index’ – 2 behaviours). Adding uninformative behaviours reduces accuracy dramatically

### Does unbalanced collection effort bias indices?

At times, it is not possible to maintain a balanced data collection effort across individuals, as not all individuals are easily observable/available or because studies demand a focus on some individuals over others. Unbalanced sampling effort did not impact the accuracy of the indices – with possible except of the DDSI, but the impact was weak (Figure 10). The precision of dyadic values did differ between balanced and imbalanced sampling efforts, but this was dependent on the index – the CSI was less precise for unbalanced data, while there was some evidence that the CRI was more precise for the unbalanced dataset (Figure 11). However, note that all individuals in the group were followed at some point, just not with the same regularity, and that dyadic interaction probabilities were held constant throughout the dataset – this is likely different from sampling schemes in real-world field sites.

**Figure 10:**
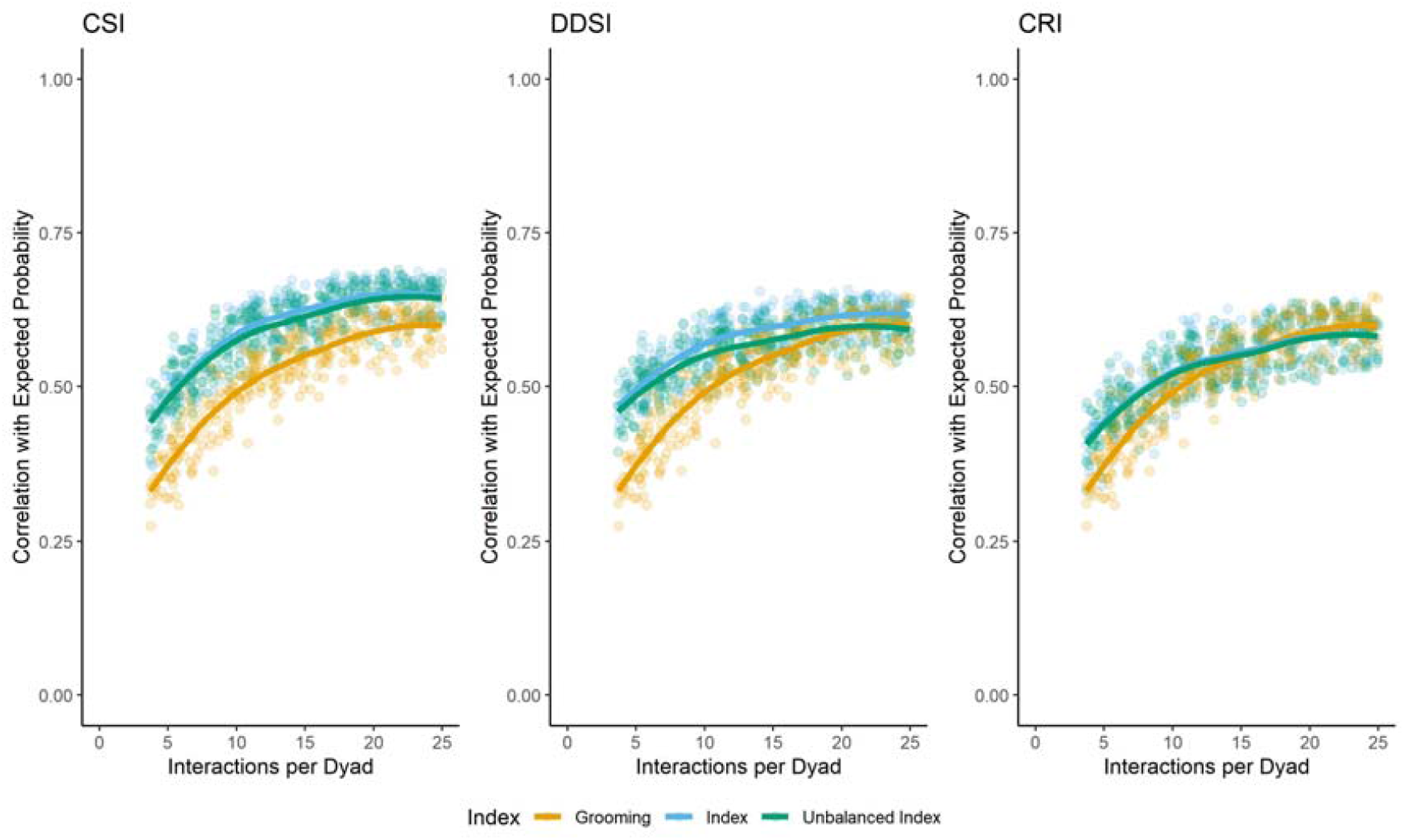
Pearson correlation between indices and the expected probability. Comparison of accuracy for the three indices with and without unbalanced focal sampling. Interaction rate of Behaviour 1/’Grooming’ shown for comparison.

**Figure 11:**
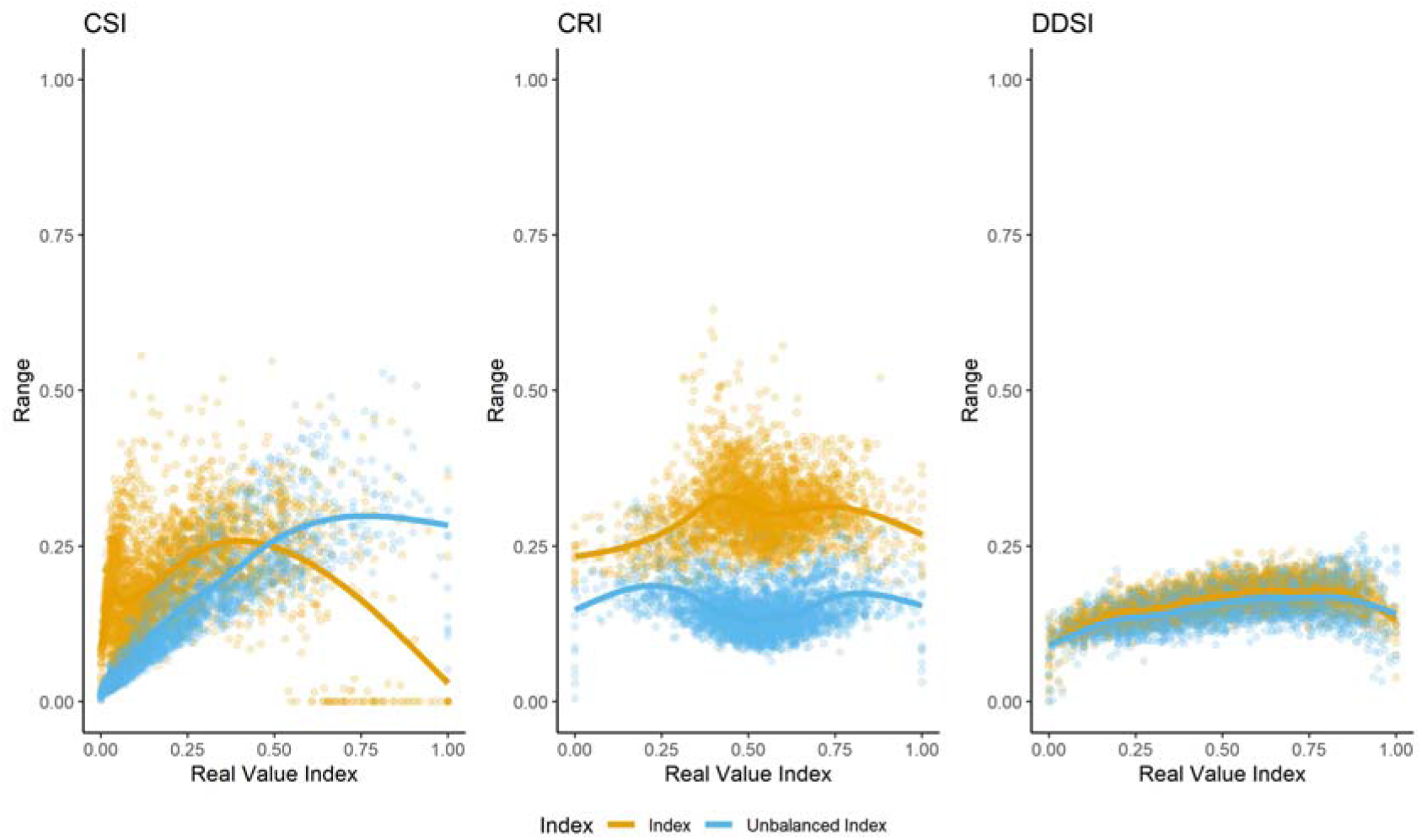
Precision for the three relationship indices with balanced and unbalanced focal sampling scheme, using only results for 360 observation days. The x-axis depicts dyadic values, standardised between 0 and 1 (i.e., 1 is the dyad with the highest value in the group). The y-axis depicts the 80%-interquantile range for 100 randomized sampling regimes. Higher range indicates lower precision.

## Discussion

In this study, we use simulations of social interactions within an animal group to determine the precision and accuracy of social relationship indices as scientific tools. Both are relevant: accuracy tells us how well an index could potentially measure what it claims to measure – that is, how similar are the observed values to the ‘true’ relationship of a dyad? Precision informs the error due to sampling bias: how likely is it that observer sampling decisions influence the estimation of dyadic values? What we show in this study is that while all three relationship indices can improve accuracy over the use of single interaction rates under specific circumstances, this is not always the case. This and their lack of improvement of precision means their overall value as instruments might be more limited than their current use suggests.

Before interpreting our results, it is important to re-iterate that while the simulated data here resemble real-world animal interaction data, they differ in important ways. The most salient is that different interaction types were based on one underlying ‘relationship’ dimension and are therefore highly correlated. This is an assumption of relationship indices but not a given in actual animals (Bergman & Beehner, 2015). The simulations also assume that dyads had stable interaction probabilities that did not vary over time. This last point means that errors of interaction rates are independent from each other, which is likely not the case in real-world data: non-random processes dictate the availability of partners for interactions, but also interaction rates on any given day. If we for example focal follow a chimpanzee on a day where the group has an intergroup encounter, we would overestimate their cooperative rates with other group members (Samuni et al., 2020). We have previously shown that this non-independence leads to low consistency in interaction rates especially for rare behaviours, conditional on sampling choices (Mielke et al., 2021). Our simulations did not include in seasonal effects, changes in underlying relationships, changes in reproductive states of females, and many other aspects that make real data interesting. Thus, the simulations might paint a more positive picture than what we would expect for a real social group.

One point to consider throughout is that we got different results for the accuracy and the precision: even for relatively small datasets, accuracy (the representation of the underlying dimension) of all indices was fairly good. However, the precision shows that dyadic values, especially for small datasets, were highly volatile. The reason for this difference is that, in right-skewed distributions, the dyads with low interaction frequencies will be low independent of sampling scheme, so the correlation between the expected and observed values will be high. However, whether a dyad was assigned a value in the highest 30% of the distribution or the lowest 30% can vary dramatically depending on who was sampled when. As we usually have one sample (our observational data), we do not know whether our data properly reflect the underlying dyadic values.

Overall, the three indices shared some advantages but also suffered the same problems. With the indices, it was faster to collect enough data to estimate the underlying relationship dimension - conditional on the fact that there is one *relationship dimension and that all included interaction types are accurate representations of this dimension*. The CSI and DDSI performed better at this than the CRI – with the latter probably performing worse because aggression, which is given higher weight in the CRI, was not as strongly correlated with the underlying dimension as the other interaction types. Given that both aggression rates and socio-positive interaction rates are usually highly right-skewed in animal groups, it is hard to imagine a linear dimension that results in both distributions if aggression is simply a mirror image of affiliation. Including aggression as a negative aspect on the same relationship dimension in the CRI and DDSI reduces precision when values are standardised (e.g., for analysis in a linear model), because dyads who never interact with each other are not on one end of the distribution, but in the centre. On average, when many aggressive events are observed in the group, dyads who never interact with each other appear to be more valuable than if there are few aggression events, but this might not reflect biology. Thus, it is worth considering whether aggression events in a species provide enough information to justify including them in one-dimensional indices.

Another aspect of all indices (and all interaction rates) was that their accuracy and precision increased with increasing sample sizes. We showed this effect previously for the consistency of real-world data (Mielke et al., 2021). The precision of indices seemed to be driven by the precision of the rarest interaction type – here, the precision of the CSI was most similar to the rarest and most imprecise interaction type, ‘food sharing’. *Social relationship indices thus do not alleviate the need for having sufficient data available per dyad per interaction type to estimate dyadic values for each of the components*. It is still common for social animal studies to calculate interaction rates and sociality indices for very short time periods with little information available for each individual, or to include extremely rare interaction types. This is particularly problematic in large groups, where inherently more data is needed to achieve adequate accuracy and precision of social indices. It is also still surprisingly rare for authors who use relationship indices to present information on the sample size for each of the components included in the index– how many grooming bouts, proximity events, etc. were involved in making the index? This is relevant, because the quality of the index and therefore the interpretability of the results can only be judged conditional on the fact that sufficient data are available. Interaction rates and indices that are based on fewer interactions than there are dyads in a group cannot possibly represent social relationships or networks accurately (Mielke et al., 2021). There is a responsibility for reviewers and editors to ask authors to make this information publicly available.

Another aspect of social relationship indices that ties into this is that unbalanced data collection did not have a strong effect on the accuracy of the indices, and had mixed results for the precision. A central tenet of field ethology is that focal follow data collection procedures produce the most accurate representation of individual social relationships, because they control for inter-individual differences in observation effort (Altmann, 1974). However, sampling imbalance affected precision and accuracy less than insufficient interaction data. If the accuracy and precision of an index or interaction rate is influenced mainly by the sample size, then focal follows are poorly suited to maximise accuracy and precision, because so many observed interactions are ignored by the observer. Scan sampling or all occurrence and ad *libitum* sampling (Altmann, 1974) could potentially circumvent this problem of lost data, but this would need a rethinking of how to calculate interaction rates and design studies using these observation methods (Canteloup et al., 2020). With new technological solutions and electronic data collection methods, scan samples and ad libitum data collection are becoming simpler to implement (Smith & Pinter-Wollman, 2020; Van Der Marel et al., 2021).

With increasing sample size, precision of relationship indices increased for the extremes – non-interacting dyads and very strong ‘friends’ were detected reliably independent of sampling decisions (conditional on dyadic interaction probabilities being expressed the same way across interaction types). However, there was low precision for dyads who interact regularly, but not every day. This is similar to precision problems reported for association indices (Whitehead, 2008). As it is common in studies of social bonds to delineate a subset of dyads (e.g., the top 10% or top 3 per individual) as ‘friends’ and assign them special significance (Silk et al., 2010a), this level of imprecision is worrisome. The values for some of these dyads will not be robust – a second observer who followed the same group on different days would assign different ‘friends’. Depending on when individuals were followed and how much data were available, results might be highly conditional on sampling biases, and we do not currently know how this affects interpretation.

Any scholar trying to make a relationship index for their study group will be faced with the question which interaction types to include in the index. Is it better to include more interaction types, even if few data are available or when the connection between the interaction type and the underlying relationship quality is unclear (e.g., infant handling, affiliative gestures)? Our results showed that the accuracy and precision of the relationship index was dependent on the accuracy and precision of the most imprecise component that made up the index. The inclusion of uninformative interaction types severely limited the ability of the index to detect the underlying relationship dimension, even if such a dimension existed. When adding uninformative or imprecise interaction rates, researchers mainly add noise. If an interaction type exists for which sufficient data is available, and which is believed to closely represent the underlying relationship (e.g., grooming), then this interaction rate might be better for most studies than just compiling interaction types together in an index. One reason for representing social relationship on a single dimension in the past was that it was easier to run univariate statistical models this way – however, linear models make this process increasingly unnecessary, and we could show that including interaction types as independent fixed effects in a linear model retained more information than combining them into an index. Thus, including independent rates into analyses of outcomes of social relationships might be more useful for most studies.

Attempts to represent social relationships or ‘bonds’ on one dimension are increasingly challenged by non-linear and multi-dimensional approaches using cluster analysis (Fischer et al., 2017) or mixture models (Weiss et al., 2019) to identify underlying relationship ‘types’ (Bergman & Beehner, 2015). Multilevel networks will also become more popular as they become easier to implement (Lehmann & Ross, 2011). However, these approaches are as sensitive to sampling biases and measurement error as relationship indices and results will be highly dependent on the interaction types researchers choose to include in their analyses. The current problem is that we do not have a clear quality measure that would differentiate a ‘good’ index/cluster solution from a ‘bad’ one. The fact that we can create an index and it has a certain distribution does not inform us about data-generating process – we cannot quantify sampling and measurement error, and how they influence the observed rates.

Social relationships in animals, under these circumstance, can best be understood as a state space model (Nielsen, 2019). Relationships cannot be observed directly, but have to be inferred after two stochastic processes take place: each interaction choice by an individual is an imperfect representation of their true relationships, and our attempts to aggregate across many interactions adds more noise. However, there should be an assignment of dyadic values that best approximates the underlying structure, given the data. It should be possible to approximate this value using probability-based approaches and likelihood maximisation, rather than relying on interaction rates. One important step to reduce at least the impact of sampling biases would be to move away from working with aggregated interaction rates as outcome variables for the study of social interactions. If we calculate dyadic values over one year of data collection, we cannot control for all the different factors that potentially influence who chooses which interaction partner at any point in time. Studying social interactions as decision situations (Kajokaite et al., 2019; Mielke et al., 2018; Samuni et al., 2018) might be more informative because more situational factors can be accounted for statistically. The ‘optimal’ relationship representation (whether one-dimensional or relationship type) would be one that optimises partner choice probabilities across social situations and interaction types, while controlling for the social environment – this is an optimisation problem that should be solvable.

What this study shows is that we need more detailed meta-scientific studies of the tools that animal behaviour researchers are using on a regular basis. Currently, researcher degrees of freedom in this area are large, resulting in known threats to replicability and reproducibility (Wicherts et al., 2016). We need more detailed and realistic simulation studies that explore the underlying properties of indices we use; however, we also need further observational studies that compare the precision and accuracy of different interaction rates and social indices (Canteloup et al., 2020; Davis et al., 2018). Importantly, researcher need to be more transparent about their choices, why they make them, and how choices influence interpretation of social outcomes – it is not enough to simply calculate interaction rates and social relationship indices because it is easily done, but researchers must show that their results are not conditional on the choices made (Asendorpf et al., 2013). One solution to this is increased transparency: readers need to know how many interactions were used to create an interaction rate, because the validity of results will be conditional on this fact. Another solution is multiverse analysis (Steegen et al., 2016): instead of calculating one relationship index and basing interpretations on this choice, researchers could simply run models based on different indices and interaction rates – if results are robust, it is unlikely that any particular researcher choice influenced analyses. However, this does not alleviate one very central problem: if insufficient data are available to estimate interaction rates, and many dyads are assigned false zeros, no measure of robustness or multiverse analysis can correctly estimate the relationship values. This study did not address all choices that researchers might face when calculating relationship indices – for example, how robust are indices to changes in underlying relationships? Should interaction rates be standardised by the group mean, and should they be standardised by the observation time or time spent in association? How should we weigh different interaction types against each other? Is it better to have study group specific solutions or use only interaction types that are generally available for all primates? This study is only a first step in a process of exploring the properties of social relationship indices and how they relate to the social lives of our study animals.

## Data Availability

R scripts for simulation: https://github.com/AlexMielke1988/Social_Relationship_Simulations. We encourage researchers to test other scenarios, but please be advised that running the simulations might take a long time.

## Acknowledgements

A.M. was funded by a British Academy Newton International Fellowship.

## Supplementary Information

### Social Relationship Indices

#### Dyadic Composite Sociality Index

The most prominent social relationship index used in animal behaviour research is the Dyadic Composite Sociality Index (CSI or sometimes DSI; Silk et al., 2003), a modification of the Composite Sociality Index (Sapolsky et al., 1997). The CSI evaluates how rates of dyadic socio-positive interactions (e.g., grooming, proximity) differ from the mean population rates, providing a relative measure indicating how dyadic relationships compare. The index typically incorporates several socio-positive interaction types that are supposed to be highly correlated (Silk et al., 2013). As such, an underlying assumption of the CSI is that the relative contribution of each interaction type to the composite measure is dependent on their variance, with highly variable interaction types (large dispersion from the population mean) more strongly influencing the index. However, variability could be a result of measurement error, so interaction types with high variation, due to insufficient data, potentially bias the index disproportionally. The CSI standardizes interactions compared to the mean of the group; thus, dyadic values are dependent on accurate data for all individuals in the group. The index has a mean of 1 and is usually right-skewed, with most dyads having low values and few having high values (see Fig. S1). In this study, we use the following formula to calculate the CSI:

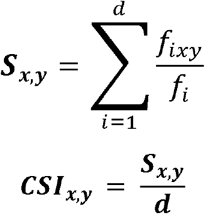

*d* – number of behaviours included in the index

*f*_*ixy*_ – rate of behaviour i for dyad xy

*f*_*i*_ – mean rate of behaviour i across all dyads in the dataset

*S*_*x,y*_ – Sum of all standardised interaction rates

We calculated a series of CSIs that included either 1) all socio-positive behaviours, that is Behaviour 1-3 (‘Grooming’, ‘Food Sharing’, ‘Proximity’); 2) socio-positive behaviours of medium or high frequency (i.e., Behaviours 1 and 3; ‘DSI common’), or 3) socio-positive behaviours of medium or high partner choice certainty (i.e., Behaviours 1 and 2; ‘DSI Interactions’). We calculated the CSI using the balanced and unbalanced datasets, and calculated a ‘bad’ version of each CSI where Behaviour 1 is replaced by an interaction distribution that is not correlated with the underlying expected probability distribution.

#### Composite Relationship Index

Considering that the entirety of dyadic interactions underlies social relationships (Hinde, 1976), including socio-positive and -negative interactions may be more informative in relationship characterization. Following this rationale, another social relationship index, the Composite Relationship Index (CRI; Crockford et al., 2012), was developed as a derivative of the CSI. Like the CSI, this index incorporates rates of different interaction types and their deviation from the population mean, with each interaction type influencing the index relative to its variance. Unlike the CSI, the CRI consists of both affiliative and agonistic interaction rates, some of which are not correlated. Doing so, the CRI can take on negative (mainly aggressive) and positive values (mainly cooperative); however, it comes with the assumption that aggression is an expression or antecedent of a ‘negative’ social relationship and on the same dimension as the socio-positive interaction types. The index also differentiates specifically between rare and common socio-positive behaviours; requiring the user to make decisions about classifying interaction types as rare/common or strongly/weakly predictive of ‘social bonds’. The CRI is roughly normally distributed around 0 (see Fig. S1). The CRI has mainly been applied in great ape research, including studies in chimpanzees (Crockford et al., 2013) and bonobos (). We used the following formula to calculate the CRI:

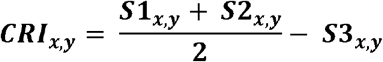

*S1* - rates of frequent socio-positive behaviours (grooming and proximity)

*S2* - rates of rare socio-positive behaviours (food sharing)

*S3* - rates of socio-negative behaviours

We calculated the CRI with all four interaction types. We also created it for both balanced and unbalanced datasets. Additionally, we created a ‘bad’ CRIs in which Behaviour 1 was replaced by a socio-positive behaviour of similar properties but uncorrelated with the underlying interaction probability.

#### Dynamic Dyadic Sociality Index

Both the DSI and the CRI provide a single averaged parameter per dyad that is fixed per observation period, thus overlooking the dynamic and flexible nature of social interactions (Wittig et al., 2020). For example, rates of interactions are subjected to changes in individuals’ reproductive or social status (Gumert, 2007), the ecological environment (Brent et al., 2013), and fluctuations in within- and between-group competition (Samuni et al., 2020), aspects that may vary over short periods of time. Thus, the Dynamic Dyadic Sociality Index (DDSI; Kulik, 2015) incorporates the dynamic aspect of social relationships into their characterization. The DDSI evaluates dyadic social relationships by applying a method similar to Elo ratings (Albers & De Vries, 2001), in which every interaction sequentially changes the dyadic value either positively (affiliative interaction) or negatively (agonistic interaction). The weight of the dyadic social change for each interaction type is determined by its frequency in the dataset, such that rarer interactions impact the index stronger. Changes in the value of a dyad lead to opposite changes in all other dyads containing the two individuals, so that the group average value remains at stable at 0.5, leading to roughly normal distributions. An advantage of the DDSI over other established methods is that it provides a daily representation of social relationships that is based on experience, thereby avoiding the assumption that relationships are fixed and can be reliably quantified by aggregation over time (Mielke et al., 2017). The DDSI has thus far been applied in studies of chimpanzees (Mielke et al., 2018; Preis et al., 2018; Samuni et al., 2018), sooty mangabeys (Mielke et al., 2018), and Assamese macaques (De Moor et al., 2020).

We calculated the DDSI using the script provided by Kulik (2015). All four behaviours were entered and weighted by their frequency of occurrence – thus, the more common an interaction type, the less each single instance affects the index. The index is updated daily and thus provides daily values; as there are no changes in the underlying dyadic probabilities, we used the DDSI for the last day of the study period to represent the whole study period, as that is the data point that includes the largest amount of information.

**Figure S1:**
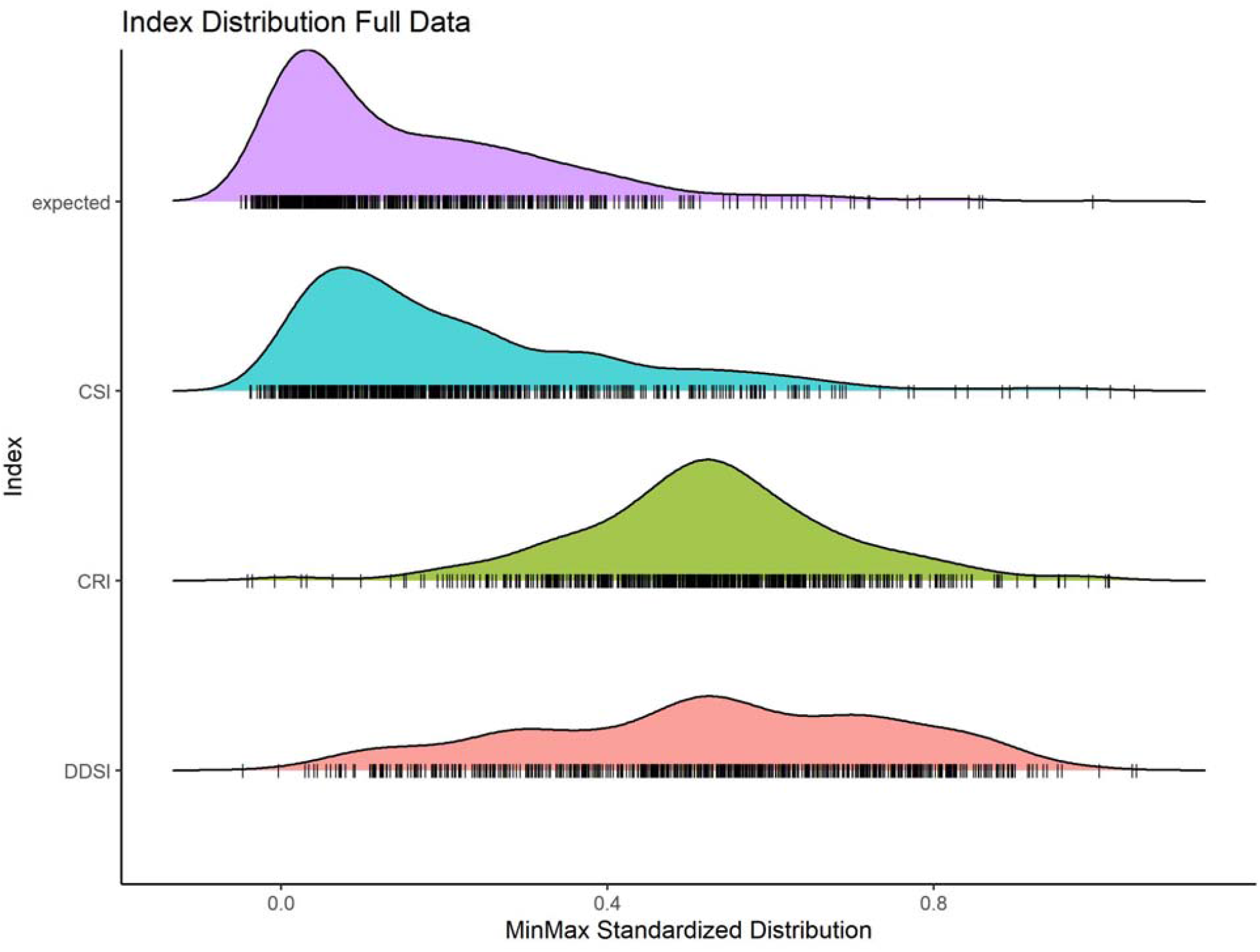
Example of distribution of the three relationship indices based on a full simulated dataset. All indices were min-max standardized between 0 and 1; naturally, the CSI has a mean of 1, the CRI of 0, and the DDSI of 0.5.

### Simulations

To assess the robustness of the different social relationship indices we simulated social interactions between individuals within different social groups of 25 individuals (10 datasets in total) over a one-year period. Each dyad was randomly assigned an expected interaction probability from a right-skewed beta distribution with α = 0.5 and β = 2. The interaction probabilities of A to B and B to A are therefore based on the same dyadic value – however, all probabilities were subsequently standardised to sum to 1 within individuals, so that there are light differences depending on direction. The final distribution differs between simulations, but the coefficient of variation for all expected probabilities was usually around 1.0, with a right-skewed distribution. Dyads with high expected probabilities interact positively a lot, while those with low expected probabilities would hardly interact with each other. As not all types of social interactions occur at the same frequency, and since different positive interaction types likely vary in how accurately they represent the value of the social relationship, we varied the frequency and certainty of occurrence of interaction types. The frequency value of the behaviour represents the average number of interactions of that type exhibited per day. The certainty of the behaviour signifies how selective partner choice is for the respective interaction type, such that a behaviour of low certainty/selectiveness has low explanatory power of relationship value and vice versa. If certainty is high, only a small number of partners are ever chosen, and the interaction distribution is highly right-skewed; if certainty is low, the likelihood of each partner is roughly even.

Socio-positive interaction types are variations on these expected dyadic interaction probabilities, but the distribution was changed by raising the probability to different powers to make them more or less certain (Behaviour 1/Grooming: 1.2; Behaviour 2/Food Sharing: 3; Behaviour 3/Proximity: 0.5). The higher the power, the larger the difference between ‘favourite’ partners and non-partners; thus, the more the favourite partners would be chosen. Interaction types also differed in how common they were: Behaviour 1 occurred 3 times per day per individual, Behaviour 2 occurred 1 time a day per individual, Behaviour 3 occurred 8 times a day per individual. Behaviours 2 and 3 were considered events, while each occasion of Behaviour 1 was assigned a duration, with dyads with higher interaction probability also having longer interactions (similar to real-world grooming).

Socio-negative interaction probabilities were calculated by taking the expected interaction probability for each dyad and taking the square root of that value 4 times and subtract the result from 1 (to switch the skew), and then again standardising probabilities to sum to 1 within individuals. Behaviour 4 was created by taking the result and raising it to the power of 2.5 to achieve the same distribution as Behaviour 1; Behaviour 4 also had 3 occurrences per individual per day, to make it the mirror image of Behaviour 1. The ‘Aggression’ distribution correlated at around r = −0.3 with the expected interaction probability and the socio-positive behaviours, similar to what we observe in some real-world social systems (see Fig. S2).

**Figure S2.**
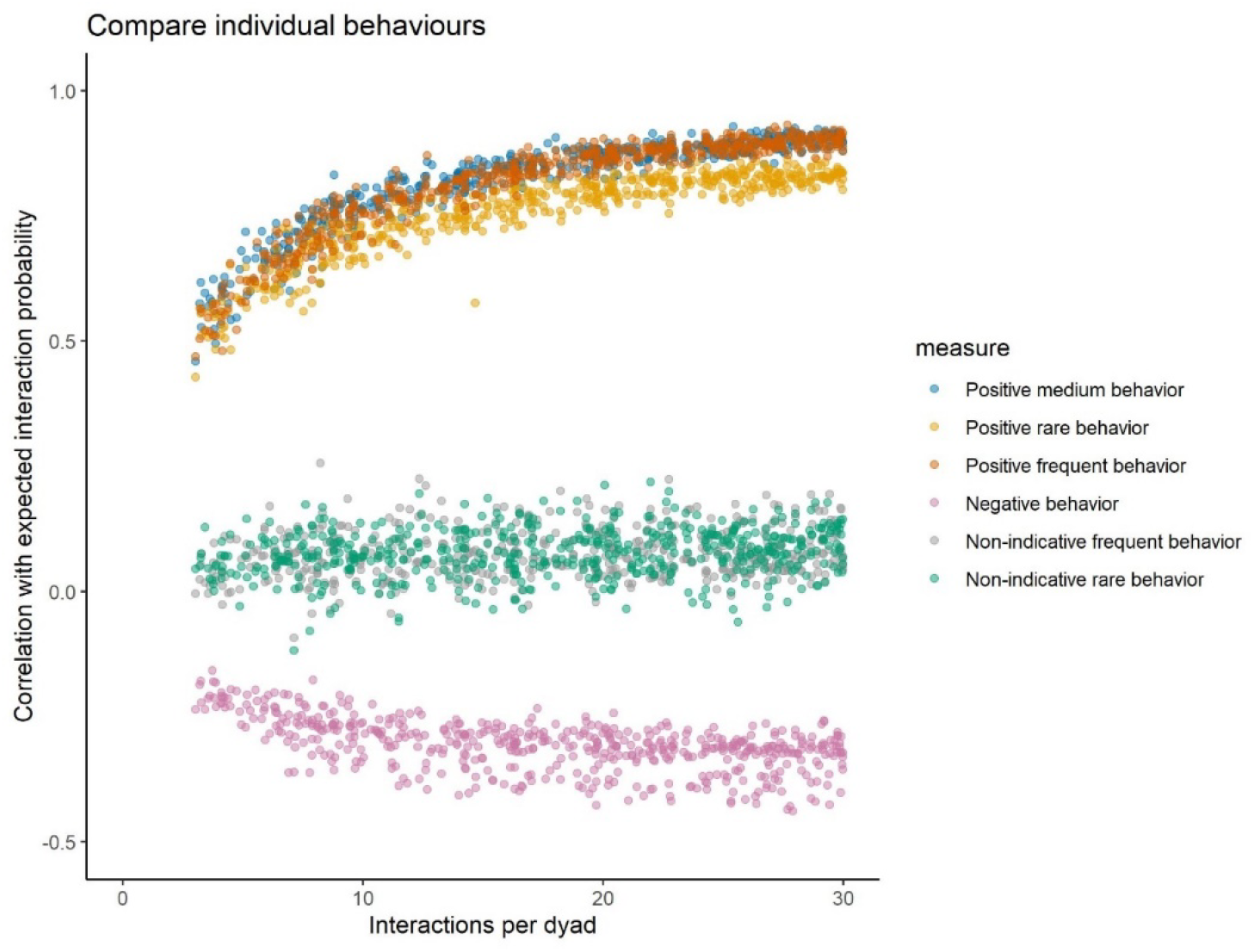
The correlation of different behaviour types with expected probabilities of interaction

For each individual for each day, a fixed number of interactions of each type was set. We introduced random variation in interaction partner choice by limiting the number of group members available as partners for any interaction. We achieved this by simulating random party compositions in which on average 50% of the group was available at any given time. Therefore, both the expected probabilities and partner availability defined who would interact in case interactions occur. For each individual, we generated a daily dataset of social interactions during 12 observation hours.

To simulate actual data collection efforts in a primate group, we selected one focal per day and varied how many days a year observers were following groups. As the overall interaction probabilities are the same for every single day, randomly selecting observation days is the same as simulating a shorter but consistent study period. Please note that this is not the same for real-world data, where seasonal effects and changes in relationships will influence results. For balanced datasets, the probability of each focal being followed was the same; for unbalanced dataset, individuals were assigned one of three weights (rare, medium, common) to indicate their probability of being chosen as a focal – in many primate field sites, one sex is followed more regularly than others, and often some individuals are harder to observe than others. For each dataset, we ran 100 iterations that randomly selected between 30 and 360 focal follow days and calculated all indices and results for each of these iterations.

Overall, we generated an ‘optimal’ dataset in which every group member was followed daily with all their interactions marked. Thus, for one year of interactions, we know who interacted how with whom. As a measure of accuracy, we report the correlations between the ‘true’ relationship and each index, and the index calculated over the full dataset and each index. As a measure of precision, we compared the dyadic values achieved across all 100 simulations. As a measure of how well each index explain the underlying partner choice probabilities, we fitted a beta regression of each index against the expected interaction probabilities and compared the Akaike Information Criterion of each model.

## Notes

https://github.com/AlexMielke1988/Social_Relationship_Simulations

### Competing Interest Statement

The authors have declared no competing interest.

